# Medium-Chain Fatty Acid exposure in non-transformed mammary glands leads to pro-tumorigenic alterations associated with aging

**DOI:** 10.1101/2025.10.01.679891

**Authors:** Mariana Bustamante Eduardo, Abul B.M.M.K. Islam, Curtis W. McCloskey, Maria Paula Zappia, Maxim V. Frolov, Rama Khokha, Elizaveta V. Benevolenskaya, Seema A. Khan, Susan E. Clare

**Author notes:** **Corresponding author**: Susan Clare.

## Abstract

The local breast environment is a valuable research focus for identifying the etiological and biological factors that contribute to the development of breast cancer. Single-cell RNA sequencing shows that the increased availability of the medium-chain fatty acid octanoic acid induces changes typical of the aged mammary gland, including downregulation of cell-cell junctions, altered extracellular matrix (ECM)-receptor interactions, and upregulation of aging markers like MDK and GDF15. *Ex vivo* exposure to medium-chain fatty acids compromises cell-cell junctions leading to cell dissemination. As aging is associated with an increased risk of developing breast cancer, it is crucial to identify the factors driving these age-related changes in the mammary gland to develop effective cancer prevention strategies. Given that the proportion of breast adipocytes increases with age, we propose that the remodeling of the mammary gland associated with aging is partly due to the increase in adipocytes and fatty-acid release.

## Introduction

Age is the major determinant of cancer risk,^1^ including breast cancer.^2^ The incidence of breast cancer increases with age although there is a slowing down of the increase starting around age 50;^3^ the probability of developing breast cancer is higher in women over the age of 50, making it the second leading cause of cancer related deaths in this age group.^4^

Aging and cancer share common hallmarks, such as genomic instability, chronic inflammation, intestinal dysbiosis, and epigenetic alterations, all of which contribute to malignant transformation and progression.^5,6^ Great efforts have been expended to fill the knowledge gap regarding how age-related changes impact the biology of the normal mammary gland and its connection to malignant transformation. Yan and collaborators,^7^ Li and collaborators,^8^ and Angarola and collaborators,^9^ have dissected at the single-cell level, the alterations associated with aging in the normal mammary gland. Additionally, Sayaman and collaborators have studied differences between luminal epithelial and basal cells from younger and older women.^10^ These comprehensive studies have identified age-related cellular and molecular perturbations in the mammary gland including loss of cell identity with age,^7,9^ transcriptomic changes,^7–9^ epigenetic changes,^9^ aberrant proliferation of epithelial cells,^7^ alterations disrupted cell-cell communication,^8^ and a proinflammatory microenvironment.^8^

The factors contributing to age-related changes in the breast are yet to be fully identified, as various factors, including hormonal changes,^2^ influence these alterations. Given that aging increases vulnerability to breast cancer, identifying non-stochastic factors that induce these changes could help develop cancer prevention strategies. The mammary gland experiences changes during aging, such as a decrease of terminal duct lobular units, decreased hormone concentrations, cellular transformation and an increased proportion of adipose tissue.^11^ Curiously, our *in vitro* and *ex vivo* data on fatty acid exposure in non-transformed mammary cells and normal tissue-derived breast microstructures show many similarities to the aged mammary gland. Based on this, we propose that the remodeling of the mammary gland associated with aging is partly due to the increase in adipocytes and fatty acid release as our data suggests that fatty acid exposure induces changes seen in the aged mammary gland, including epigenetic changes,^12^ elevated ROS,^12^ downregulation of cell-cell junctions, altered extracellular matrix (ECM)-receptor interactions, and increased expression of aging-related genes such as GDF15 and MDK which may be pro-tumorigenic (Figure 1A).^7,8^

**Figure 1.**
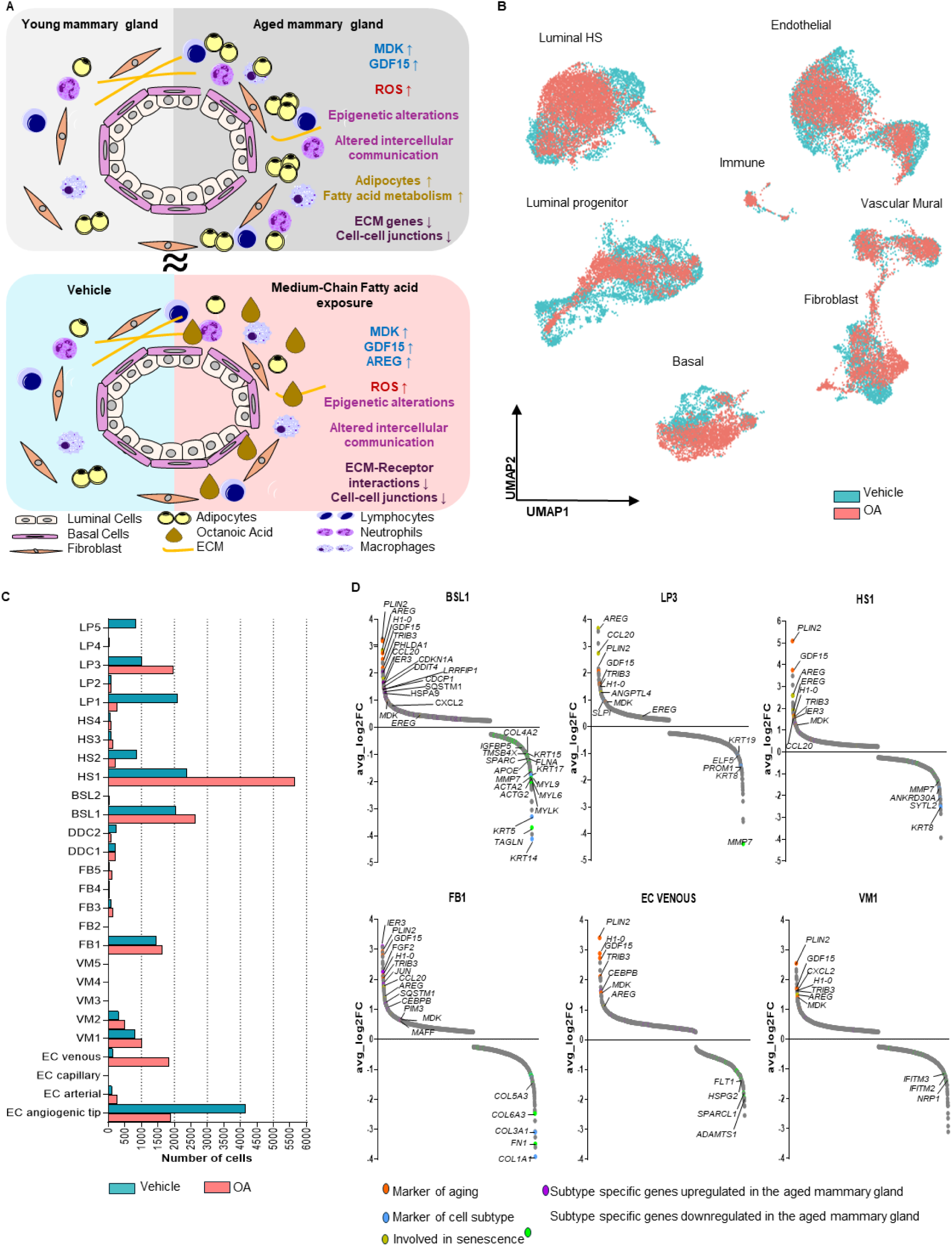
Octanoic Acid (OA) exposure induces cell composition and transcriptional changes resembling aged mammary tissue. **(A)** Medium-chain fatty acid exposure causes changes typical of the aged mammary gland. **(B)** Approximation and Projection (UMAP) depicting the major mammary cell types. Plot colored by treatment based on scRNA-seq data. Luminal hormone sensing (HS), luminal progenitor, basal, endothelial, immune, vascular mural, fibroblast. **(C)** Bar plot showing the number of cells within each subtype in vehicle- or OA-treated breast microstructures, based on scRNA-seq data (N = 2). Cell subtypes defined by Reed et al.^60^ **(D)** Differentially expressed genes (adj p < 0.01) upon octanoic acid (OA) exposure compared to vehicle in epithelial and non-epithelial cell subtypes. Genes are aligned along the *x*-axis according to average log2 fold change (FC). Positive values are up regulated with OA and negative values are downregulated with OA.

## Results

### Exposure to medium-chain fatty acids induces age-related gene expression changes and loss of cell identity markers in epithelial and non-epithelial mammary cells

We have used *in vitro* and *ex vivo* models to study the effects of the medium-chain fatty acid octanoic acid (OA) in non-transformed breast epithelial cells and in breast tissue.^12,13^ When compared to vehicle, changes induced by OA such as increases in ROS, changes in epigenetics, and cell-cell communication landscape alterations, resemble those reported for the aged mammary gland. To further explore the connection between OA exposure and aging, we analyzed the single cell RNA sequencing (scRNAseq) data set (GSE284557) that we generated to explore the effects of OA in tissue-derived breast microstructures.^12^ OA alters the distribution of cell clusters of epithelial and non-epithelial mammary cell populations (Figure 1B). In the epithelial compartment, we identified epithelial mammary cell populations including basal (BSL, 1-2), hormone sensing (HS, 1-4), LPs (1-5) cells with as well as cells that show a mix of basal and HS markers referred as donor derived clusters (DDC, 1 and 2). In the stromal compartment, we identified fibroblast (FB,1-5), vascular mural (VM, 1-5) and endothelial cells (EC, venous, capillary, arterial and angiogenic tip). Cell proportions were affected by OA exposure, notably the fraction of BSL1, HS1, LP3, FB1, VM1, VM2 and EC venous cells increased upon exposure (Figure 1C).^12^

We observed significant (p < 0.01) gene expression changes induced by OA; notably, OA significantly upregulates aging-related genes in both epithelial and non-epithelial compartments such as midkine (MDK),^7^ Growth Differentiation Factor 15 (GDF15),^14,15^ Perilipin 2 (PLIN2),^16^ Tribbles Pseudokinase 3 (TRIB3),^17^ Chemokine Ligand 2 (CXCL2),^18^ and H1.0 Linker Histone (H1-0)^19^ (Figure 1D). One of the downregulated genes upon OA exposure was MMP7, a gene whose reduced expression contributes to the accelerated aging of mammary epithelial cells.^20^ Furthermore, OA exposure led to the significant upregulation of genes shown to have increased expression in specific subtypes within the aged mammary gland and downregulation of genes with decreased expression in specific subtypes within the aged mammary gland (Figure 1D). Lineage markers were also repressed by OA in epithelial subtypes and fibroblasts (Figure 1D). These results show that medium-chain fatty acid exposure causes gene expression changes that are typical of the aged mammary gland.

### Exposure to medium-chain fatty acids upregulates senescence related pathways

Senescence can be regarded as a key hallmark of aging.^21^ Therefore, we examined differentially expressed senescence-related genes in both epithelial and non-epithelial compartments. We observed significant (p < 0.01) upregulation of components of the Senescence-Associated Secretory Phenotype (SASP)^22^ in both epithelial and non-epithelial subtypes (Figure 1C), such as amphiregulin (*AREG*), Epiregulin (*EREG*), C-C Motif Chemokine Ligand 20 (*CCL20*) and Angiopoietin Like 4 (*ANGPTL4*). Reactome pathway analysis in the three epithelial subtypes that increase their proportion upon OA, BSL1, LP3, HS1, revealed an upregulation of terms related to senescence including SASP (Figure S1A). These results show that medium-chain fatty acid exposure induces the expression of senescence-related genes.

### Cell-cell communication analysis reveals an increase in MDK and GDF15 signaling following medium-chain fatty acid exposure

We employed cell-chat to identify key signaling pathways and interactions between different cell populations within breast microstructures affected by OA exposure. A total of 41,222 inferred interactions were identified in vehicle condition while 13,424 were identified under the OA condition, including secreted signaling, ECM-receptor interactions and cell-cell contact (Figure 2A-B and Figure S1B). Among the enriched ligand-receptors pairs from the secreted signaling category were AREG, MDK and GDF15 (Figure 2C). When analyzing BSL1, LP3, HS1, FB1, VM1 and VM2, which are the subtypes that showed an increase in proportion upon OA (Figure 1C), we found that AREG, GDF15 and MDK were among the strongest signals to their specific receptors in most epithelial, stromal, and immune cells upon OA exposure (Figure 2C and Figure S2). The receptors predicted to be interacting with MDK were SDC1, 2 and 4, PTPRZ1, ITGA4-ITGB1, ITGA6-ITGB1, LRP1, NCL and ALK. GDF15 interacted with TGFBR2. AREG interacted with EGFR and EGFR-ERBB2. Notably, the expressions of *AREG*, *GDF15* and *MDK* were induced in most cell subtypes (Figure 3A). Additionally, the cell culture supernatant from breast microstructures embedded in Matrigel showed a significant increase in GDF-15 levels after 24 hours of OA exposure, confirming the secretion of this protein (Figure S3A). These results show that the overall interactions between cell subtypes changed upon exposure to OA. Notably, we identified that among the increased secreted signaling in OA were the aging-related ligands MDK and GDF15, as well as the senescence-related ligand AREG.

**Figure 2.**
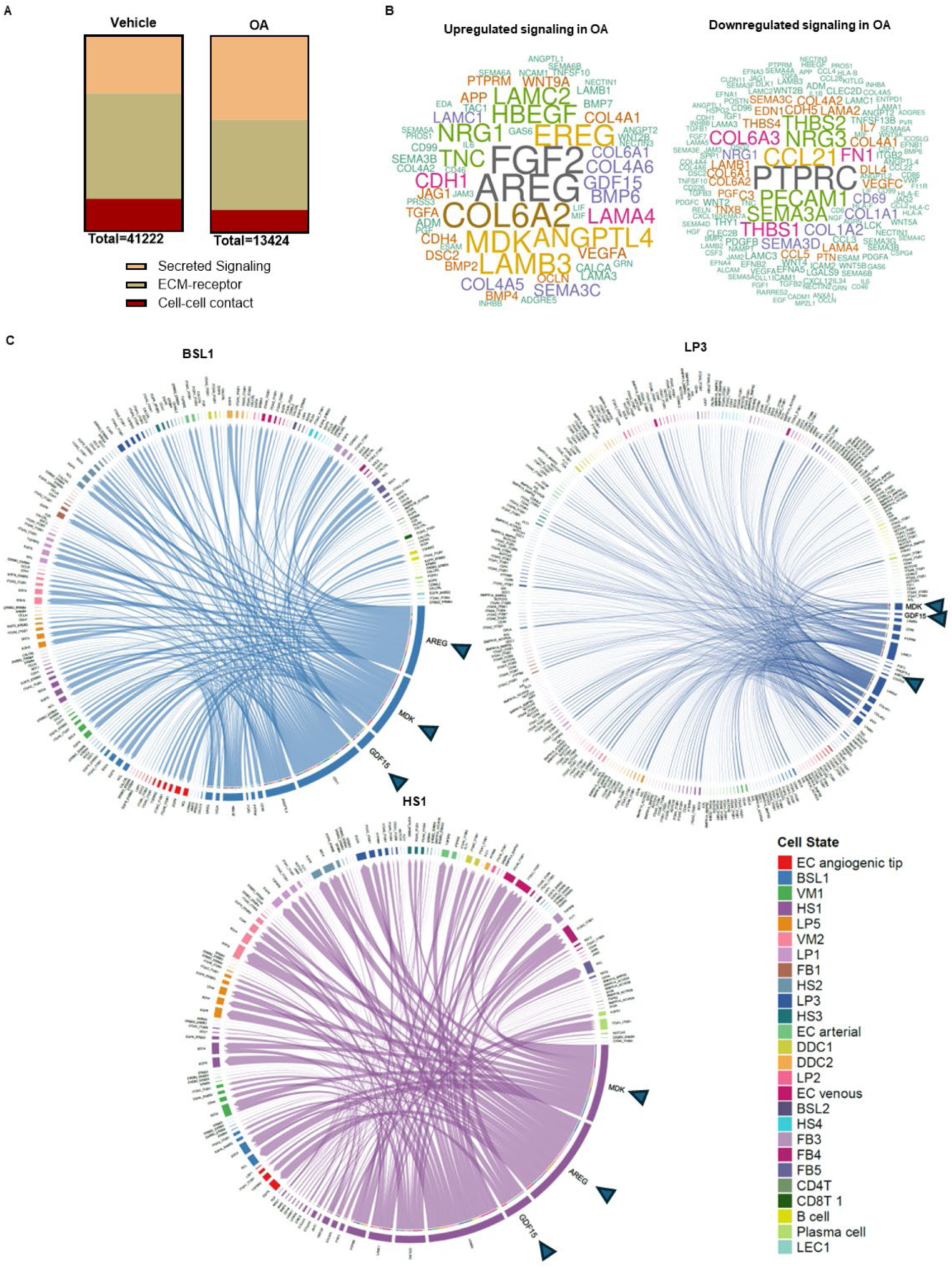
Inferred cell-cell communications patterns in mammary cells. **(A)** Number of inferred cell-cell communication in vehicle and octanoic acid (OA). **(B)** Wordcloud visualizing the up-regulated and down-regulated ligands upon OA exposure. Word size indicates the extent of enrichment. **(C)** Chord diagram for visualizing cell-cell communication of upregulated ligands in response to OA in basal 1 (BSL1), luminal progenitor 2 (LP3) and hormone sensing 1 (HS1). Arrows indicate the AREG, MDK, and GDF15 ligands.

**Figure 3.**
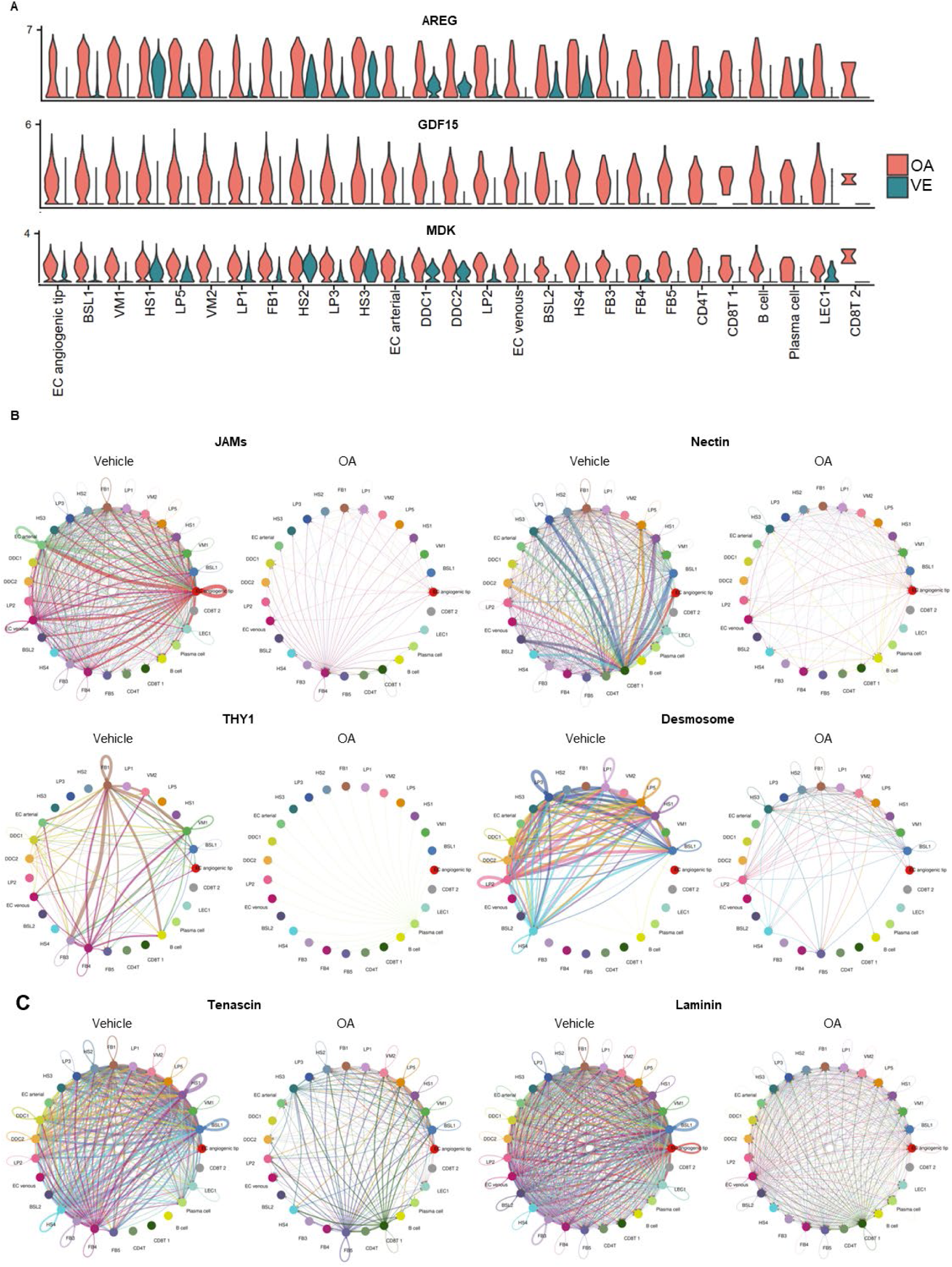
Intercellular communications analysis of mammary cells. **(A)** Violin plots displaying expression level of selected genes in vehicle and octanoic acid in epithelial and non-epithelial cell subtypes. **(B)** Circle plot showing inferred cell-cell contact pairs which decreased with OA compared to vehicle. **(C)** Circle plot showing inferred ECM-receptor interactions which decreased with OA compared to vehicle. Lines represent ligand-receptor interactions. Thickness indicates strength. Loops indicate autocrine signaling.

### Cell-cell communication analysis indicates a decrease in ECM-cell interactions and cell adhesions following medium-chain fatty acid exposure

To further examine key interactions affected by OA, we focused on the decreased interactions upon OA exposure. CellChat identified that among the interaction types present in the vehicle, most of the ECM-receptor and cell-cell contact categories are absent or decreased with OA (Figure S1B). The cell-cell contact-related pathways identified in the vehicle that decreased with OA, include the Junctional Adhesion Molecules (JAMs), the nectin family of cell-adhesion molecules, THY1, and desmosomes (Figure 3B and Figure S1B). Cell-cell adhesion pathways present in vehicle and absent upon OA exposure included: interecellular adhesion molecule (ICAM), claudin (CLDN) and CDH5 (Figure S1B). ECM-receptor interactions pathways that decreased upon OA exposure included tenascin and laminin (Figure 3C and Figure S1B). Among the ECM-receptor interaction pathways present only in the vehicle group were fibronectin 1 (FN1) and thrombospondin (THBS1) (Figure S1B). Interestingly, proteomics analysis also showed that exposure of the non-transformed mammary cell line MCF-10A to OA resulted in the downregulation of proteins associated with the ECM such as FN1, ITGAV, ITGB1, THBS1 and CDC42 (Figure S3B). These analyses revealed that cell-cell interactions and ECM-receptor interaction decreased with OA exposure.

### Medium-chain fatty acid exposure compromises basal barrier leading to cell dissemination

Given the downregulation of contractility and basement membrane genes, the reduction in cell-cell interactions, and results from the 3D acinar assay with MCF-10A cells in Matrigel, where OA treatment led to basement membrane breaches rather than enhanced cell survival or lumen filling (Figure S4), we posited that OA exposure may instead facilitate or enhance malignant cell invasion. We examined the interaction between labeled basal and luminal cell populations within breast microstructures embedded in Matrigel using confocal microscopy. Exposure to OA disrupted the architecture of the breast microstructure, allowing cells to escape their normal constraints and disseminate (Figure 4A). Similarly, 3D mammary sphere culture with primary mammary cells embedded in Matrigel showed that upon OA exposure, 3D spheres were disrupted allowing the dissemination of cells (Figure 4B). Breast microstructures and 3D spheres not only exhibit a loss of architectural integrity but also show that luminal cells are among those escaping basal constraints. Mitochondrial staining confirms cell viability, while laminin staining indicates the loss of lumen structure (Figure 4B and Figure S5). <u>These</u> analyses not only validated the findings on cell-cell communication, showing a decrease in both cell-cell and ECM-receptor interactions, but also revealed that OA exposure induces an invasive phenotype, likely due to disruption of basal barrier function.

**Figure 4.**
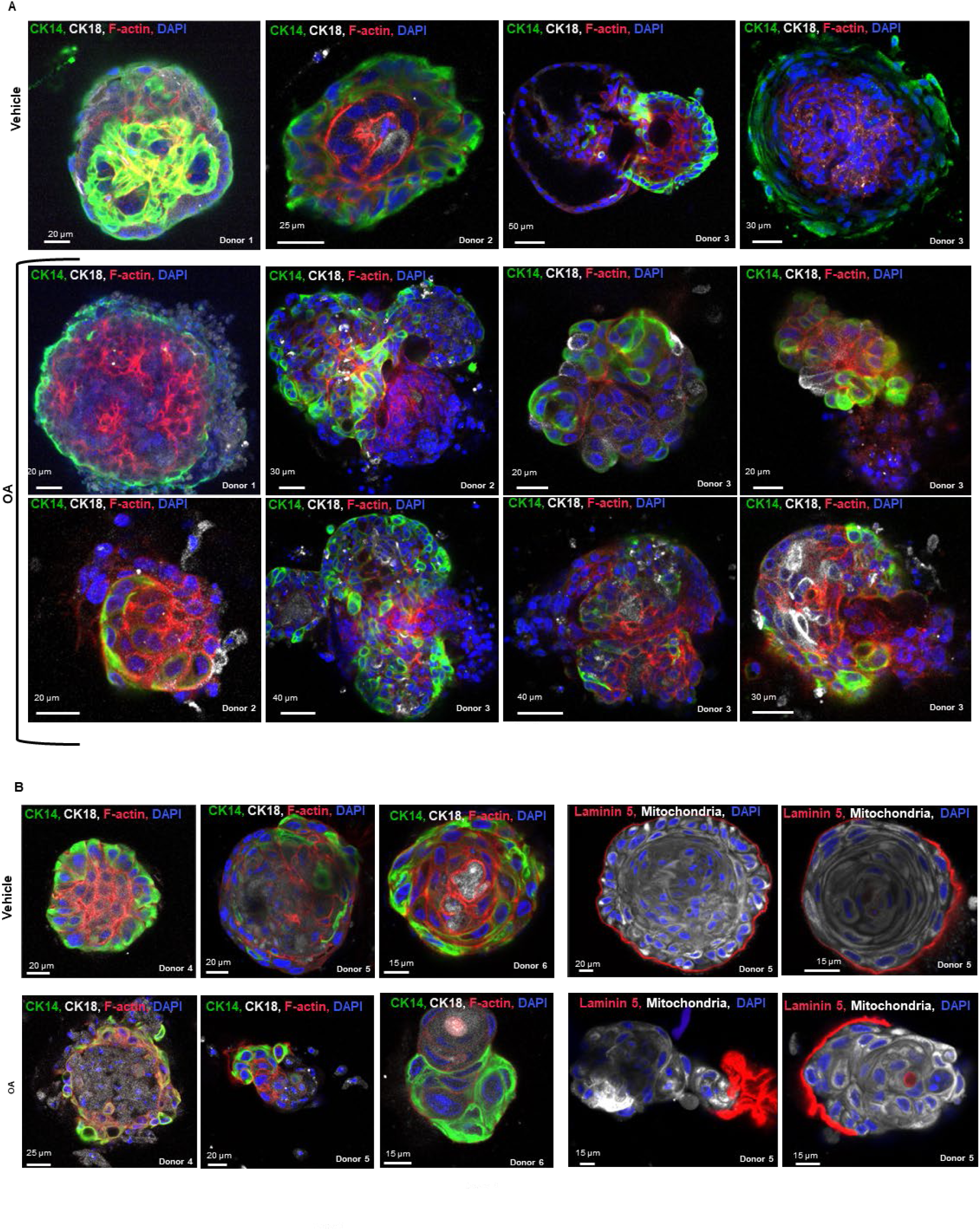
Effect of octanoic acid (OA) on breast microstructures and mammary breast spheres. **(A)** Representative fluorescence confocal sections of breast microstructures embedded in Matrigel culture in vehicle of OA containing media for 7 days. CK14=Cytokeratin 14, CK18 = Cytokeratin 18. **(B)** Representative fluorescence confocal sections of breast spheres in Matrigel culture in vehicle of OA containing media for 7 days.

### Media containing medium-chain fatty acids selects cells expressing GDF15, MDK, AREG

Seven-day OA exposure of breast microstructures and 3D spheres embedded in Matrigel reveal architecture disruption and cell dissemination. To determine which cell types are selected in OA-containing media, we adapted a previously established method for generating primary breast cell cultures as a selection strategy.^23,24^ Under these conditions, epithelial cell migration and microstructure disruption differed between vehicle and OA media, mirroring observations in the 3D models (Figure 5A). We performed scRNAseq of migrating cells and identified 16 clusters (Figure 5B), revealing a shift in cell proportions, specifically, a reduction in fibroblasts, and increased LP3, BSL1, and an uncharacterized *AREG^+^*/*GDF15^+^* population (Figure 5C). *GDF15, AREG, MDK*, and *PLIN2* were identified as general markers of migratory cells in OA-containing media. These analyses suggest that the LP3, BSL1, and AREG⁺/GDF15⁺ cell populations are best equipped to survive in medium-chain lipid–rich media.

**Figure 5.**
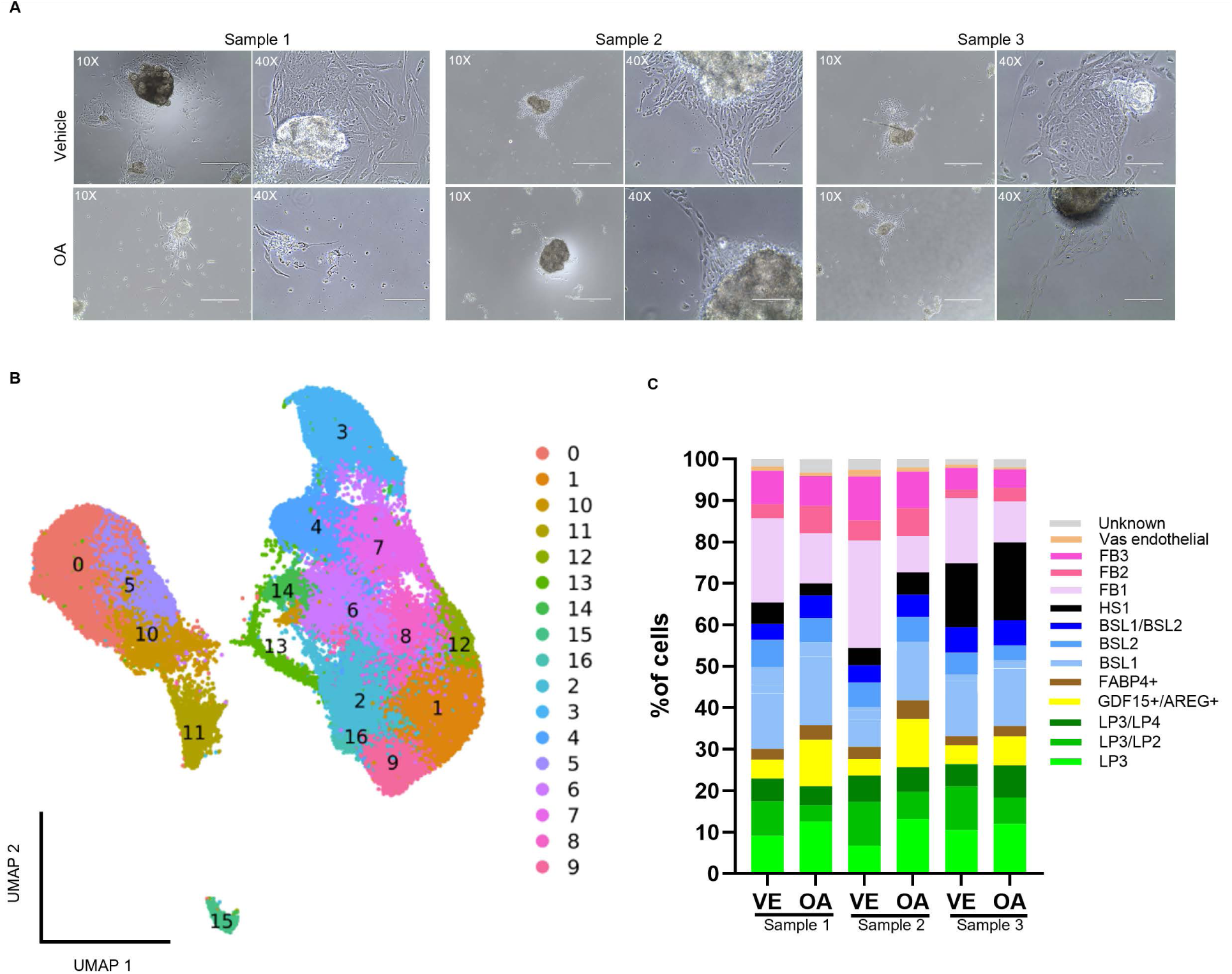
Single cell RNA analysis of cells that migrate from microstructures in vehicle of OA medium. **(A)** Images of three sequenced samples display cells migrating from breast microstructures in either vehicle or OA-containing media after 48 hours. **(B)** UMAP plot colored by cluster identity. Clusters: 0 = FB1, 1 = BSL1, 2 = LP3, 3 = HS1, 4 = LP2/3, 5 = FB3, 6 = GDF15⁺/AREG⁺, 7 = LP3/4, 8 = BSL2, 9 = BSL1/2, 10 = FB2, 11 = FABP4⁺, 12 = BSL1, 13 = BSL1, 14 = Unknown, 15 = Vascular endothelial, 16 = Unknown **(C)** Comparison of cell proportions between vehicle and OA for all three samples.

## Discussion

Previous work from our group has focused on defining the in-breast environment to identify factors that promote estrogen receptor negative breast cancer and that may be disrupted for prevention and has identified a lipid metabolism gene signature associated with the risk of this subtype. ^25,26^ To uncover the mechanistic link we studied medium-chain fatty acids effects on non-transformed breast cells *in vitro* and on tissue derived microstructures *ex vivo and* showed that exposure to OA results in metabolic dysregulations, epigenetic fostered phenotypic plasticity, cell-cell communication landscape changes and increases in ROS. ^12,13^ Surprisingly, many OA-induced changes resemble those reported in the aged mammary gland. Therefore, in this study we analyzed the scRNA-seq dataset of tissue-derived breast microstructures exposed to either vehicle or OA. We show that medium-chain fatty acid exposure causes changes typical of the aged mammary gland, such as downregulation of cell-cell junctions and ECM-receptor interactions, and transcriptomic changes identified in the aged mammary gland. Our data suggests that exposure to medium-chain fatty acids mimics changes observed in the aged mammary gland. This raises the question: what similarities exist between our *in vitro/ex vivo* study and the aged mammary gland? The answer seems to lie in the increased availability of fatty acids, as the proportion of adipose tissue in the breast increases with age.^11^ The connection between lipids stored in the breast (e.g. adipocyte-derived fatty acids from triglycerides) and breast cancer has been demonstrated.^27–29^ We propose that the fatty acids in the immediate breast microenvironment influence mammary cells similarly to our *in vitro/ex vivo* exposure, suggesting that the remodeling of the mammary gland associated with aging is partly driven by the increase in adipocytes and fatty acid release.

OA modulated gene expression in both epithelial and non-epithelial compartments, inducing the expression of aging-related genes, while repressing lineage markers and genes known to be downregulated in aging. Altered expression of lineage markers has been reported as a consequence of aging causing a reduced ability to maintain lineage fidelity and increase cell plasticity.^7,9,10^ *GDF15* is among the aging-related genes induced by OA, and its release by breast microstructures increased upon OA exposure. It is not only a marker of aging,^14^ but also a promising biomarker for various cancers,^30–32^ including breast cancer,^33^ where its levels correlate with estrogen receptor negative breast cancer.^34^ GDF15 also promotes epithelial-to-mesenchymal transition (EMT), invasiveness, the acquisition of stem cell properties, induces lipolysis and enhances fatty-acid oxidation.^34–38^ Increases in ROS due to enhanced fatty acid oxidation may explain the upregulation of GDF15, as ROS activate stress-responsive genes like the transcription factors ATF3 and ATF4, which subsequently upregulate GDF15.^39^ GDF15 has been shown to induce lipolysis in adipocytes, which results in leukemia cell growth.^40^ GDF15 may contribute to the age-dependent rise in breast tumorigenesis by its role in lipolysis. We propose that by inducing lipolysis in adipocytes, GDF15 releases free fatty acids that signal epithelial and non-epithelial mammary cells similar to our *in vitro* and e*x vivo* exposure to OA, driving fatty acid oxidation, increasing ROS, altering metabolism, enhancing fatty acid availability, and sustaining the cycle, which would facilitate oncogenic transformation. MDK was also induced by OA, we have shown that MDK expression increases upon OA exposure as a result of epigenetic reprogramming, which occurs due to the metabolic shift toward the serine pathway induced by medium-chain fatty acids.^12^ MDK is a key player in breast cancer progression,^41^ and has been recognized as a mediator of the age-associated rise in breast tumorigenesis. ^7^

Among the OA induced genes were also genes involved in senescence pathways. Senescence can be considered a central hallmark of aging, with opposing roles in cancer.^21^ On one hand, it can act as a tumor suppressor,^42^ but on the other hand, strong evidence suggests that through the SASP, it contributes to tumor progression,^43^ stemness,^44^ and EMT.^45^ Genes induced by OA are components of the SASP and have been described to promote reprogramming (AREG)^46^, cancer progression (EREG)^47^, EMT (CCL20)^48^ and migration ( ANGPTL4)^49^.

We have shown that OA leads to the decreased of cell-cell junctions and ECM-receptor interactions, something that is also observed in the aged mammary gland,^8^ and we also have shown that the consequences of the decrease cell-cell junctions and ECM-receptor interactions enhance malignant cell invasion by impairing basal cells’ ability to prevent luminal cell dissemination. Sirka and collaborators propose that basal cells are a dynamic barrier to epithelial cell dissemination and it disruption leads to cell invasion and dissemination.^50^ Interestingly, adipocyte-breast cancer cell co-culture studies have shown that lipolysis releases free fatty acids, which sustain invasion.^51^ Additionally, GDF15-overexpressing breast cancer cells exhibit a significant increase in invasion through the basement membrane matrix.^34^ Our scRNAseq analysis of cells migrating in medium-chain fatty-acid rich media reveals populations of metabolically agile, disseminating cells characterized by the expression of *GDF15, AREG, MDK* and *PLIN2*. *GDF15* promotes EMT, invasion, stem cell properties and lipolysis,^34–38,40^ *AREG is associated with* reprogramming, proliferation, invasion and motility,^46,52,53^ *MDK* promotes progression, neurogenesis, proliferation and migration,^41,54^ and *PLIN2,* a lipid droplet associated protein, has been shown to enhances EMT.^55,56^

Taken together, we propose that the release of free fatty acids from an increased number of breast adipocytes, potentially due to elevated GDF15-induced lipolysis during aging, is a factor that induces mammary gland remodeling, increasing vulnerability to breast cancer. This opens up preventive posibilities, such as targeting GDF15 itself, as antibodies against it are already being tested in clinical trials for cancer cachexia and as adjuncts to immunotherapeutic agents.^57,58^

We acknowledge that further studies are needed to demonstrate the parallels between our in *vitro/ex vivo* data and the aged mammary gland. Future studies will include lipidomic and transcriptomic analyses of both young and aged human mammary glands.

## Material and Methods

### ScRNA-seq data analysis

ScRNA-seq data from breast microstructures obtained from two donors, treated with either vehicle (PBS) or OA, corresponding to dataset GSE284557, were analyzed as described.^12^ SingleR package (version 2.0.0)^59^ was utilized for cell annotation according to the Reed *et al.* reference dataset.^60^

### Single Cell Pathway Analysis

Pathway analysis was performed using SCPA (1.2.0) with msigdbr v7.5.1 Reactome pathways,^61^ as previously described. ^12^ Senescence related Reactome pathways were selected for visualization.

### Cell-cell communication network analysis

Interactions between cell subpopulations were explored using the CellChat R package (v1.6.1.60).^62^ Secreted signaling, cell–cell adhesion signaling, and ECM–receptor signaling were assessed by evaluating the expression of ligand–receptor pairs across cell types in vehicle and OA conditions, using the CellChat R toolkit.

### Mammary microstructures preparation

Tissues from women admitted for reduction mammoplasty recruited under an approved IRB protocol (NU15B07) were collected. All participants provided written informed consent. Breast tissue was processed as previously described,^12^ briefly Tissue was minced in a sterile Petri dish and transferred to 50 ml tubes containing 1 mg/ml collagenase from Clostridium histolyticum (Sigma Aldrich, #C0130) in Kaighn’s Modification media (Gibco, #21127022) supplemented with 0.5% BSA (Sigma, #SLCM0392) and Antibiotic-Antimycotic (Gibco, #15240062), the collagenase-containing medium was filtered through a 0.22 μm filter prior to use. Falcon tubes were sealed and shaken overnight (16 h) at 100 rpm and 37 °C for gentle tissue dissociation. The following day, microstructures were collected by centrifugation (114 × g, 5 min), washed with PBS, and resuspended in HPLM medium (Gibco, #A4899101) supplemented with H14 additives and Antibiotic-Antimycotic (Gibco, #15240096), then transferred to ultra-low attachment six-well plates (Corning, #CLS3471).

### 3D culture of breast microstructures

Breast microstructures were resuspended in Matrigel (Corning, # 354230) as described by Sirka et al.^50^ with minor modifications. Matrigel was polymerized at 37°C for at least 30 minutes prior the addition of microstructures. Breast microstructures were resuspended in complete HPLM medium containing 2 % Matrigel plated in 8-well chamber slides (Thermo Scientific, # 154534). The next day, the media was change to complete HPLM containing 2 % Matrigel plus vehicle or complete HPLM containing 2 % Matrigel plus OA. Breast microstructure were incubated for one week with complete HPLM containing 2 % Matrigel plus vehicle or OA changed every 2 days.

### 3D sphere culture

Breast microstructures were dissociated into single cells as previously described.^12^ In brief, breast microstructures were incubated with pre-warmed (37 °C) TrypLE (Gibco, #12604013) for 15 minutes at 100 rpm and 37 °C, with gentle pipetting using wide-orifice P1000 tips every 5 minutes. Cell dissociation was stopped by adding pre-warmed complete HPLM media with 0.1% BSA, followed by centrifugation at 200×g for 5 minutes. Cells were then treated with 2 ml RBC lysis buffer (Invitrogen, #00433357) for 1 minute at room temperature, quenched with complete HPLM + 0.1% BSA, and centrifuged again. The pellet was resuspended in 1 ml complete HPLM containing 5 mg Dispase II (Sigma, #D4693) and 0.1 mg DNase I (Stemcell Technologies, #07469), incubated for 3 minutes at 200 rpm and 37 °C. After adding more complete HPLM, primary cells were passed 3 times through an 18G needle attached to a 10-ml syringe and filtered through a 40 µm strainer. Luminal/basal sphere cultures were performed as previously described with minor modifications.^63^ Briefly, primary cells were isolated from breast microstructures, resuspended in complete HPLM containing 2 % Matrigel and overlaid onto 8-well chamber slides pre-coated with polymerized Matrigel. Spheres formed by 14 days of culture after which cultures were treated with complete HPLM containing 2 % Matrigel plus vehicle or complete HPLM containing 2 % Matrigel plus OA for 7 additional days.

### 3D spheres and microstructures immunofluorescence staining

Whole-mount immunofluorescence staining of Matrigel-embedded spheres and microstructures was performed as previously described,^64^ using pre-warmed buffers and solutions maintained at 37°C, with minor modifications. Briefly, samples were washed once with pre-warmed PBS followed by incubation with pre-warmed 2% paraformaldehyde (PFA) at room temperature for 15 minutes. Samples were then washed 3 times with pre-warmed PBS-Glycine solution for 10 minutes each on a horizontal shaker, follow by a 20-minute incubation with pre-warmed IF-Wash buffer^64^ at room temperature. Next, samples were washed 3 times with pre-warmed IF-Wash buffer for 10 minutes each on a horizontal shaker. Samples were incubated in pre-warmed blocking solution (2% Normal goat serum in IF-Wash buffer) for 1 hour at room temperature followed by 2 days incubation at 4°C with antibody mixture. The antibody mixture included: 1:250 Cytokeratin 14 monoclonal Alexa Fluor 488 conjugated antibody (Novus Biologicals, # NBP234403X), 1:250 Cytokeratin 18 monoclonal Alexa Fluor 647 (Novus Biologicals, # NBP234414Y), Phalloidin Labeling Probes Alexa Fluor 594 (Invitrogen, # A12381), DAPI (Invitrogen, #D1306), 200 nM MitoView™ Fix 640 (Biotium, #70082), 1:250 Human Laminin alpha3/Laminin-5 Alexa Fluor® 594-conjugated Antibody (R&D Systems, Catalog # FAB2144T). After the 2 days incubation, samples were let sit at room temperature for 1 hour, then washed with pre-warmed IF-Wash buffer. After the 2 days incubation, samples were let sit at room temperature for 1 hour, then washed with pre-warmed IF-Wash buffer for 30 minutes on a horizontal shaker followed by 3 washes with pre-warmed IF-Wash buffer for 20 minutes on a horizontal shaker and 4 washes with pre-warmed PBS for for 20 minutes on a horizontal shaker. Samples were then mounted with fructose glycerol solution and stored at 4°C for 24 hours. Samples were imaged with confocal microscopy (Nikon A1R (B) GaAsP) at magnification 20 x magnification and then the images were analyzed using Imaris software.

### Single Cell analysis of migrating cells

To enrich for cells that robustly migrate in response to OA, primary breast epithelial cell cultures were established and exposed to OA-containing media under selective conditions as described.^23,24^ Breast microstructures from 3 donors were plated in Corning Primaria Cell Culture Plates (Corning, # 353801) in vehicle media (HPLM with PBS) or OA media (HPLM plus 5mM OA). Cells were allowed to migrate from the breast microstructure onto the floor of the culture dish for 48 hours as described. Cells were incubated in 0.05% Trypsin-EDTA (Gibco, # 25300054) for 15 minutes and cells were passed through a 40um strainer. Trypan blue staining was used to assess cell viability and perform cell counting. A total of 3,400 viable cells/μL were resuspended in Cell Suspension Buffer for library preparation using the Illumina Single Cell 3’ RNA Prep, T10 kit (Illumina, #20135691), following the manufacturer’s protocol. Prepared libraries were subsequently sequenced on the Illumina NovaSeq X Plus platform. Unsupervised clustering identified 17 clusters (resolution of 0.6). Cell were annotated manually according to the Reed *et al.* reference dataset.^60^

### Proteomics analysis

MCF-10A cells (ATCC, CRL-10317) cultured in HPLM medium (Gibco, #A4899101) supplemented with H14 additives were be exposed to vehicle (PBS) or 5mM OA for 24 hrs. Twenty-four hours after exposure, cells were lysed using a buffer composed of 0.5% SDS, 50 mM ammonium bicarbonate (Ambic), 50 mM sodium chloride, and a protease inhibitor cocktail (Halt, Thermo Scientific, #78430). The lysates were then sonicated on ice for three 15-second cycles. Following sonication, samples were centrifuged at 15,000 × g for 20 minutes at 4 °C to remove debris, and the resulting supernatant was collected. Protein concentration was measured using BCA and micro BCA assays (Thermo Scientific, # 23227, #23235). A 65 μg protein aliquot was precipitated overnight with cold acetone and TCA (8:1 ratio), then the pellet was resuspended in 100 μL of 8 M urea in 0.4 M ammonium bicarbonate and vortexed to mix. Proteins were reduced with 4 μL of 100 mM DTT at 55 °C for 30 minutes, then alkylated with 18 mM iodoacetamide for 30 minutes in the dark at room temperature. Urea was diluted to 1.8 M by adding four volumes of ultrapure water. Digestion was performed overnight at 37 °C using sequencing-grade trypsin (1:50 enzyme-to-protein ratio), and stopped by adding formic acid to 0.5%. Peptides were desalted with Pierce C18 spin columns (Thermo Scientific), eluted in 80% acetonitrile with 0.1% formic acid, dried by vacuum centrifugation, and reconstituted in 30 μL of 5% acetonitrile with 0.1% formic acid for LC-MS analysis. For mass spectrometry analysis, peptides were analyzed using a Dionex UltiMate 3000 nanoLC system coupled to an Orbitrap Elite mass spectrometer (Thermo Fisher Scientific). Samples were first loaded onto a 150 μm × 3 cm trap column and separated on a 75 μm × 10.5 cm PicoChip C18 column, both packed with 3 μm ReproSil-Pur® beads. Chromatography was performed at 300 nL/min using a 120-minute gradient from 5% to 40% acetonitrile in 0.1% formic acid. MS1 scans were acquired in the Orbitrap (400–2000 m/z, 60,000 resolution), with an AGC target of 1 × 10⁶. The top 15 precursor ions were fragmented by CID (35% energy) in the ion trap. Dynamic exclusion was set to 60 s. All analyses were conducted in triplicates. Raw MS/MS data were analyzed with Mascot (v2.5.1) against the SwissProt database. Searches included fixed carbamidomethylation (Cys) and variable modifications: methionine oxidation, asparagine/aspartic acid deamidation, and N-terminal acetylation, allowing up to two missed tryptic cleavages.

Peptide-spectrum matches were filtered at 1% FDR, and only proteins with at least two unique peptides were retained. Identifications were visualized and assessed using Scaffold (v5.0, Proteome Software).

### 3D acinar assay

MCF-10A cells were cultured as previously described in DMEM/F12 (Gibco, #11320033) supplemented with 5% horse serum (Gibco, #26050070), 20 ng/ml epidermal growth factor (EGF) (Gibco, # PHG0311L), 10 μg/ml insulin (Lonza, # BE02-033E20), 0.5 μg/ml hydrocortisone (STEMCELL Technologies, # 07925), 100 ng/ml cholera toxin (Sigma, # C8052, and antibiotics (Gibco, # 15240096).^65^ To generate acini, MCF-10A cells were seeded on Matrigel and grown in DMEM/F12 supplemented with 5 ng/ml EGF, 2% horse serum, 0.5 μg/ml hydrocortisone, 10 μg/ml insulin, 100 ng/ml cholera toxin, antibiotics and 2% Matrigel as previously described.^66^ Acini were cultured for 21 days, with media changed every 4 days. For antioxidant treatment, Trolox (Abcam, #ab120747) was added to both the Matrigel and the culture media. Octanoic acid was added to the media during the final 7 days of the culture period. Immunofluorescence staining of acini was carried out following established protocols, ^66^ using an Alexa Fluor 488-conjugated cleaved caspase-3 antibody (Cell Signaling Technology, #9669S) and an Alexa Fluor 594-conjugated laminin-5 antibody (Bio-Techne, #C21441T). Nuclear counterstaining was performed with DAPI (Invitrogen, #D1306). To assess luminal filling, confocal microscopy (Nikon A1R (B) GaAsP) was used to image the central region of each acinus at magnification 20 x magnification and then the images were analyzed using Imaris.

## Resource availability

Requests for further information and resources should be directed to and will be fulfilled by the lead contact, Susan E. Clare.

## Acknowledgments

We thank Robert H. Lurie Comprehensive Cancer Center Flow Cytometry Core Facility for cell sorting on a BD FACSymphony S6 SORP system, purchased through the support of NIH 1S10OD011996-01 and 1S10OD026814-01. Imaging work was performed at the Northwestern University Center for Advanced Microscopy (RRID: SCR_020996) generously supported by NCI CCSG P30 CA060553 awarded to the Robert H Lurie Comprehensive Cancer Center. Proteomics services were performed by the Northwestern Proteomics Core Facility, generously supported by NCI CCSG P30 CA060553 awarded to the Robert H Lurie Comprehensive Cancer Center, instrumentation award (S10OD025194) from NIH Office of Director, and the National Resource for Translational and Developmental Proteomics supported by P41 GM108569. We would like to thank Basil Baby Mattamana (Aarohan M) and Raju Gajjela from the Northwestern Proteomics Core for the proteomics work. This work was supported by the Northwestern University NUSeq Core Facility for scRNA-seq. We are grateful to our many lab colleagues for constructive feedback. This work was supported by the 2023 AACR-Pfizer Breast Cancer Research Fellowship, Grant Number 23-40-49-BUST (M.B.E.), the Breast Cancer Research Foundation, Avon Foundation and Bluhm Family Foundation Foundation (to S.A.K.), the National Cancer institute R21CA288676-01A1 (to S.E.C.).

## Author contributions

Conceptualization, M.B.E, S.A.K, and S.E.C.; Methodology, M.B.E., A.B.M.M.K.I., and C.W.M.; Software, .B.M.M.K.I., and C.W.M.; Validation, M.B.E.; Formal analysis, M.B.E., A.B.M.M.K.I., C.W.M., and S.E.C.; Investigation, M.B.E, and P.M.Z.; Resources, S.A.K, and S.E.C; Writing - Original Draft, M.B.E, S.A.K, and S.E.C.; Writing - Review & Editing, M.B.E, S.A.K, and S.E.C.; Visualization, M.B.E.,and A.B.M.M.K.I., Supervision, R.K., M.V.F., E.V.B., S.A.K, and S.E.C.; Funding acquisition, M.B.E, S.A.K, and S.E.C.

## Declaration of generative AI and AI-assisted technologies

No generative AI or AI-assisted technologies were used in the creation of this work.

