## Supplementary material for "Medium-Chain Fatty Acid exposure in non-transformed mammary glands leads to pro-tumorigenic alterations associated with aging": Figure S

### Supplementary Figure1

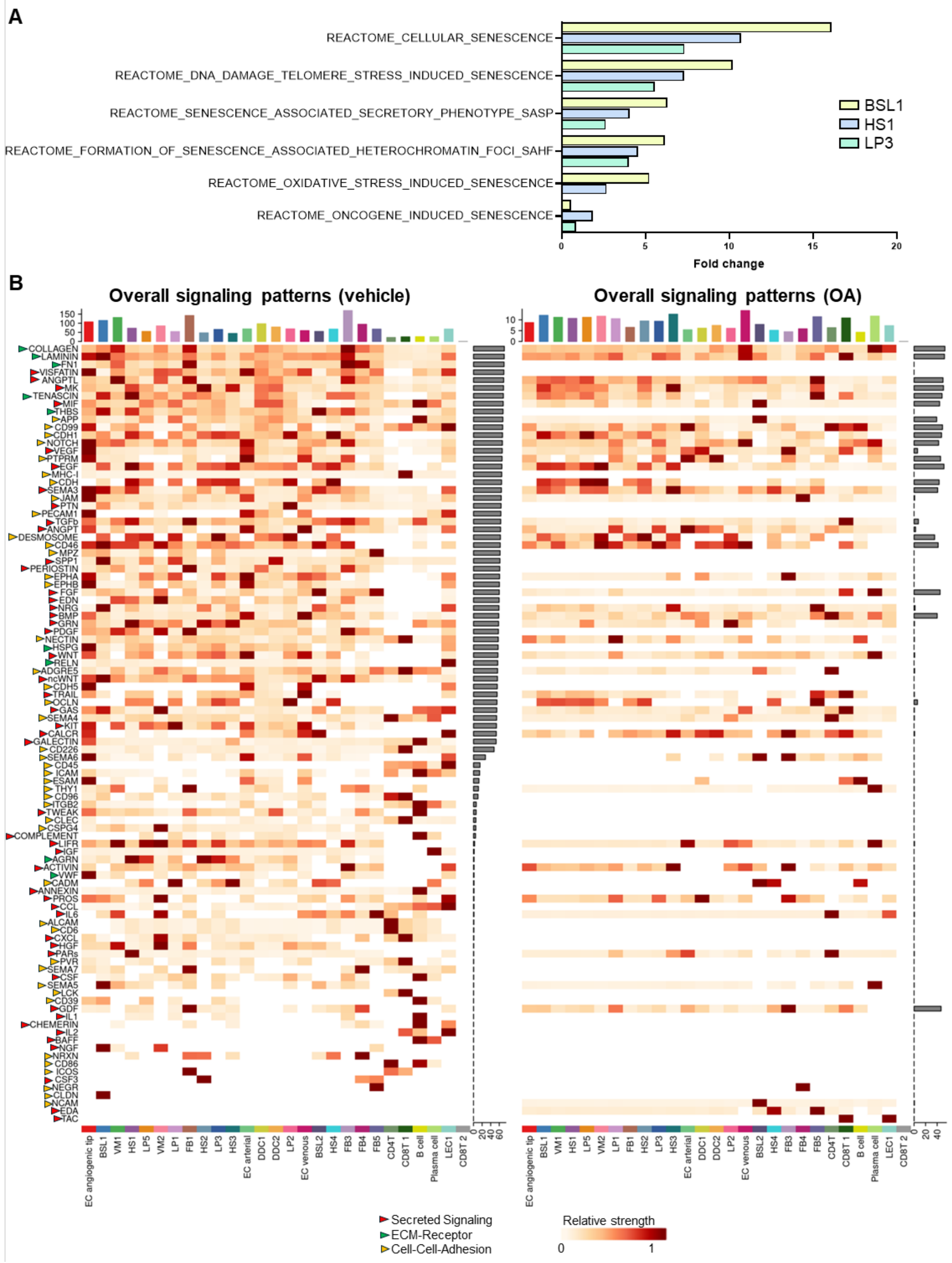

**Figure S1. Analysis of single-cell profile of tissue-derived breast microstructures treated with vehicle and octanoic acid (OA**

**(A)** Reactome analysis of upregulated senescence-dependent terms in basal 1 (BSL1), luminal progenitor 2 (LP3) and hormone sensing 1 (HS1).

**(B)** Overview of intercellular communication between vehicle and octanoic acid (OA) treated breast microstructures.

**Supplementary Figure 2**

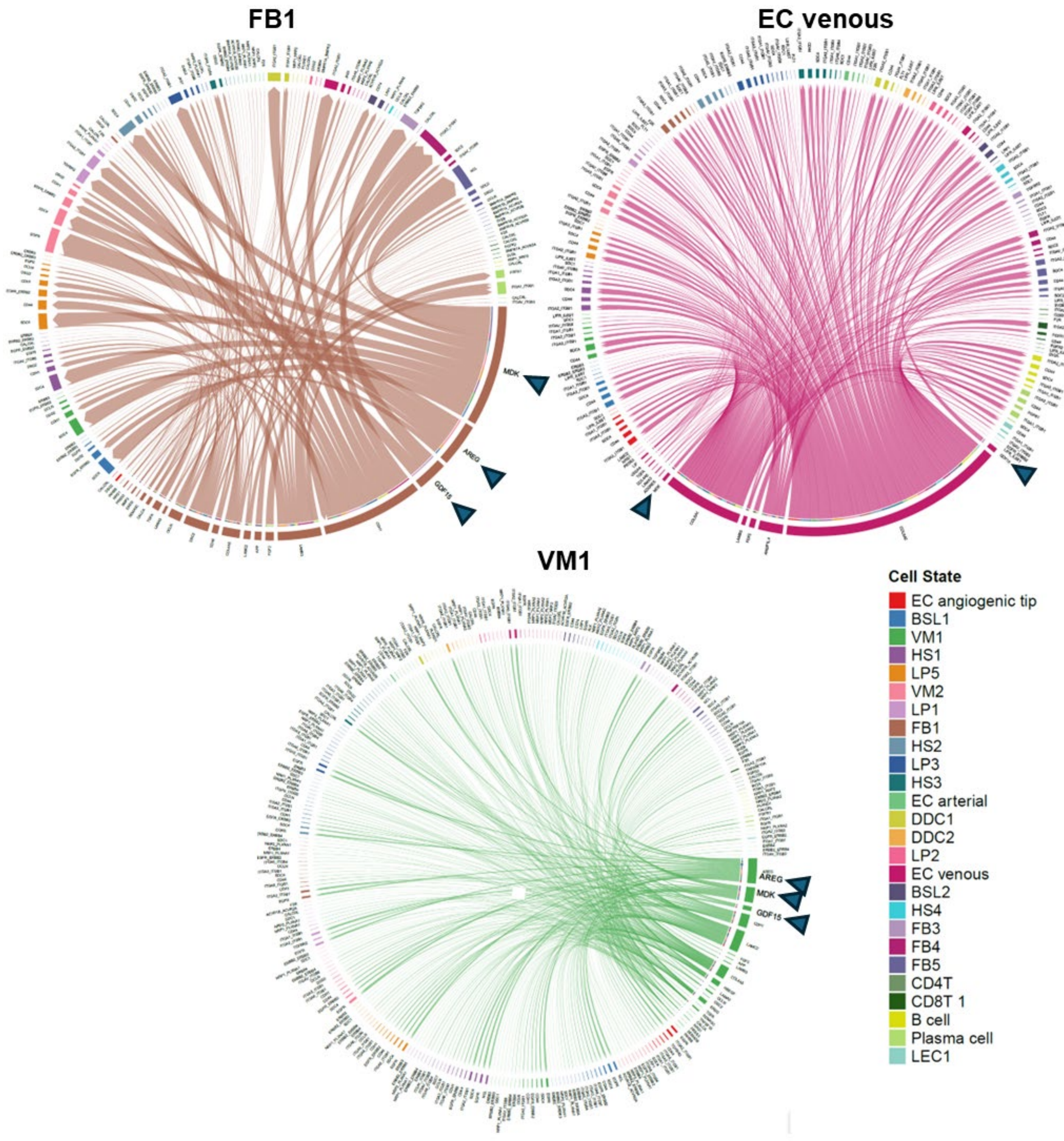


**Figure S2. Cell-cell communication of upregulated ligands upon octanoic acid (OA) exposure.** Chord diagram for visualizing cell-cell communication of upregulated ligands in response to OA in fibroblast 1 (FB1), endothelial cell (EC) venous, vascular mural 1 (VM1) and vascular mural 2 (VM2). Arrows indicate the AREG, MDK, and GDF15 ligands.

**Supplementary Figure 3**

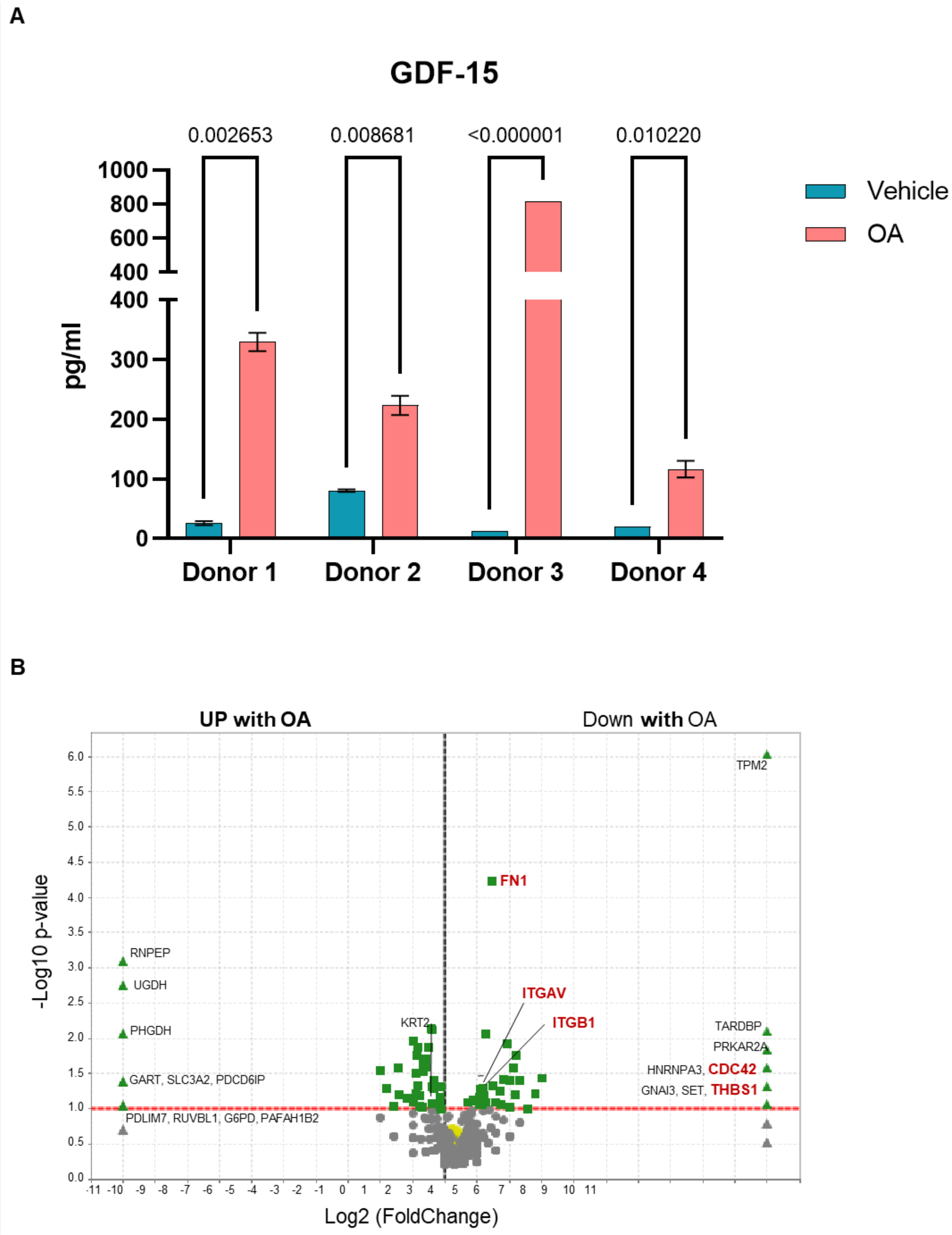

**Figure S3. Differential analysis of proteins upon octanoic acid (OA) exposure in MCF-10A cells.**

- **(A)** Quantification of GDF-15 in the supernatant of microstructures embedded in Matrigel after 24-hour exposure to vehicle (PBS) or OA, measured by ELISA. Bars represent mean ± SEM. Statistical analysis was performed using a t-test (n = 2).
- **(B)** Volcano plot shows differentially expressed proteins between OA and vehicle. Proteins involved in ECM-receptor interactions are depicted in red.

**Supplementary Figure 4**

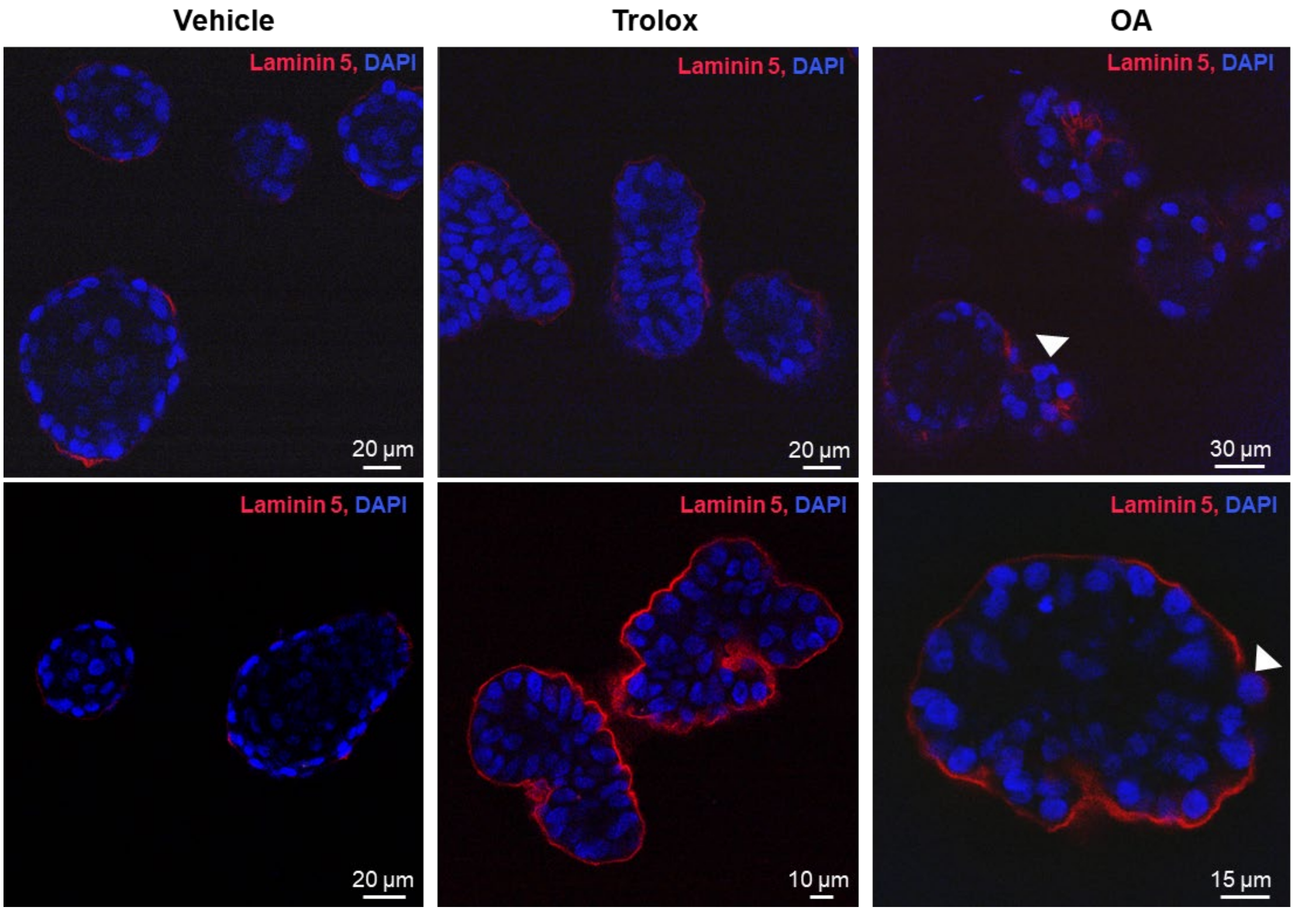


**Figure S4. 3D acinar assay.** Control (vehicle): clear luminal filling (increased ROS and decreased GSH cause cell detachment, leading to anoikis and autophagy). Trolox: cells detach and fill the lumen. OA: basement membrane breach (arrow).

**Supplementary Figure 5**

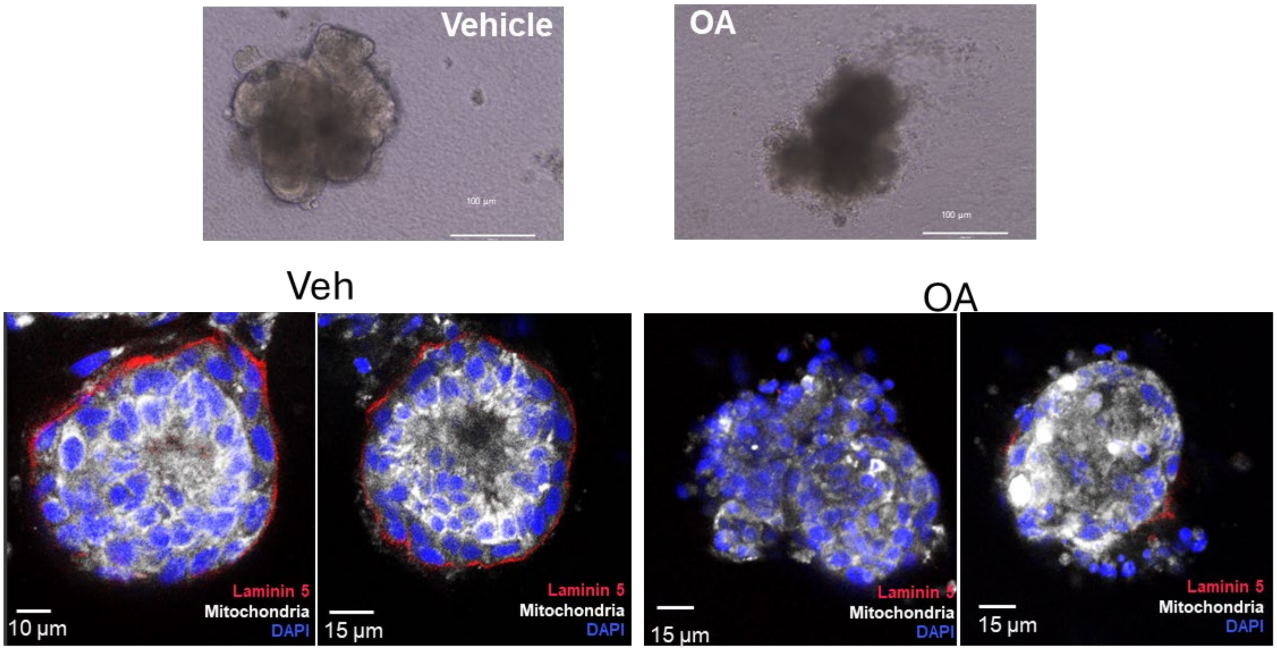


**Figure S4. Effect of octanoic acid (OA) on breast microstructures and mammary breast spheres.** Representative breast microstructures visualized with phase contrast microscopy or with confocal microscopy.
